# Interleukin 7 receptor (IL7R) is required for myeloid cell homeostasis and reconstitution by hematopoietic stem cells

**DOI:** 10.1101/2020.09.02.280362

**Authors:** Taylor Cool, Atesh Worthington, Donna Poscablo, Adeel Hussaini, E. Camilla Forsberg

## Abstract

Respiratory diseases are a leading cause of death worldwide, with highly varied vulnerability to disease between individuals. The underlying reasons of disease susceptibility are unknown, but often include a variable immune response in lungs. Recently, we identified a surprising novel role of the interleukin 7 receptor (IL7R), a primarily lymphoid-associated regulator, in fetal-specified, lung-resident macrophage development. Here, we report that traditional, hematopoietic stem cell-derived myeloid cells in the adult lung, peripheral blood, and bone marrow also depend on IL7R expression. Using single and double germline knockout models, we found that eosinophil numbers were reduced upon deletion of IL7Rα. We then employed two Cre recombinase models in lineage tracing experiments to test whether these cells developed through an IL7Rα+ pathway. Despite the impact of IL7Rα deletion, IL7R-Cre labeled only a minimal fraction of eosinophils. We therefore examined the intrinsic versus extrinsic requirement for IL7R in the production of eosinophils using reciprocal hematopoietic stem cell transplantation assays. These assays revealed that extrinsic, but not eosinophil-intrinsic, IL7R is required for eosinophil reconstitution by HSCs in the adult lung. To determine which external factors may be influencing eosinophil development and survival, we performed a cytokine array analysis between wild-type and IL7Rα-deficient mice and found several differentially regulated proteins. These findings expand upon our previous publication that IL7R is required not only for proper lymphoid cell development and homeostasis, but also for myeloid cell homeostasis in tissues.

**Highlights:**

- Loss of IL7Rα resulted in significantly fewer eosinophils in adult mice
- IL7R-Cre lineage tracing revealed minimal labeling of eosinophils
- IL7Rα-deficient HSCs robustly reconstituted eosinophils in a WT host
- WT HSCs failed to fully reconstitute eosinophils in IL7Rα^-/-^ hosts
- Several cytokines are differentially expressed in WT and IL7Rα-deficient mice

---

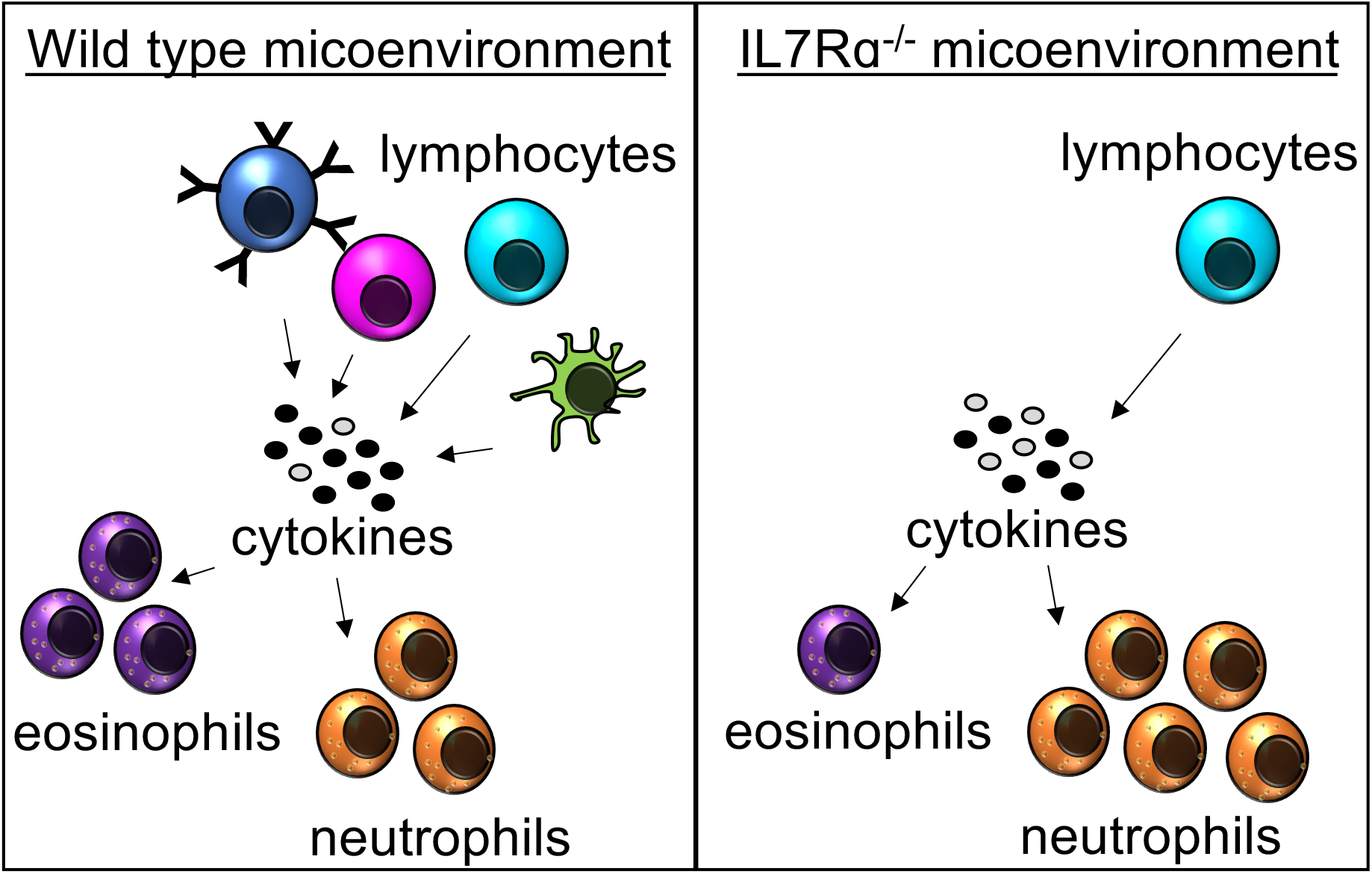

## Introduction

Hematopoiesis is the process of mature blood and immune cell production and maintenance from hematopoietic stem and progenitor cells (HSPCs). The majority of “traditional” blood and immune cells, including eosinophils, have short half-lives and are continually replaced by HSPCs^1^. In contrast, fetally-derived tissue resident macrophage (trMacs) self-maintain in their respective tissues for life, without contribution from adult HSCs, thereby leading us to consider them “non-traditional” mature immune cells^2–5^. Recently, our lab uncovered a novel role for interleukin 7 receptor (IL7R) in trMac specification during development^6^. Though IL7R had originally been reported to be essential exclusively for lymphocyte development and survival, a few recent studies, including ours, have found that it also plays roles in myeloid cell development^6–8^. Interestingly, in addition to our finding that IL7R regulates trMac specification during mouse fetal development^4^, studies in humans have reported that other myeloid cell types, including monocytes and eosinophils, may express IL7R^7,9^. Furthermore, these cells can upregulate IL7R mRNA and surface protein upon stimulation with lipopolysaccharide (LPS), a known activator of eosinophils. Here, we used germline knockout mice, lineage tracing, and transplantation assays to determine whether IL7R plays a role in adult-derived “traditional” myeloid cell types in the lung, peripheral blood (PB), and bone marrow (BM).

## Results and Discussion

### Adult neutrophils and eosinophils are differentially affected by deletion of IL7Rα

During our previous studies where we found that IL7R unexpectedly regulates trMac development, we also noted alterations in other cells types in the lungs of IL7Rα^-/-^ mice. We therefore employed Flk2^-/-^ and IL7Rα^-/-^ mice and crossed the two strains to also create homozygous double mutants (referred to as FIDKO mice here) to investigate lung cellularity in three mutant cohorts. Flk2 is a tyrosine kinase receptor that regulates hematopoietic development^10^, including robust numbers of common myeloid progenitor cells (CMPs)^10,11^, the presumed precursors of eosinophils. Previously, our lab established that all adult-derived “traditional” mature blood and immune cells, but not trMacs^6,12^, develop through a Flk2+ pathway^6,11,13,14^. As expected based on previous reports of circulating lymphocytes^15^, B lymphocytes in the lungs were decreased in all three mutant strains relative to the WT controls, with drastic reductions in the IL7Rα^-/-^ and FIDKO mice (Fig.1A), while neutrophil numbers were normal in all three genetic models (Fig. 1B).

**Figure 1:**
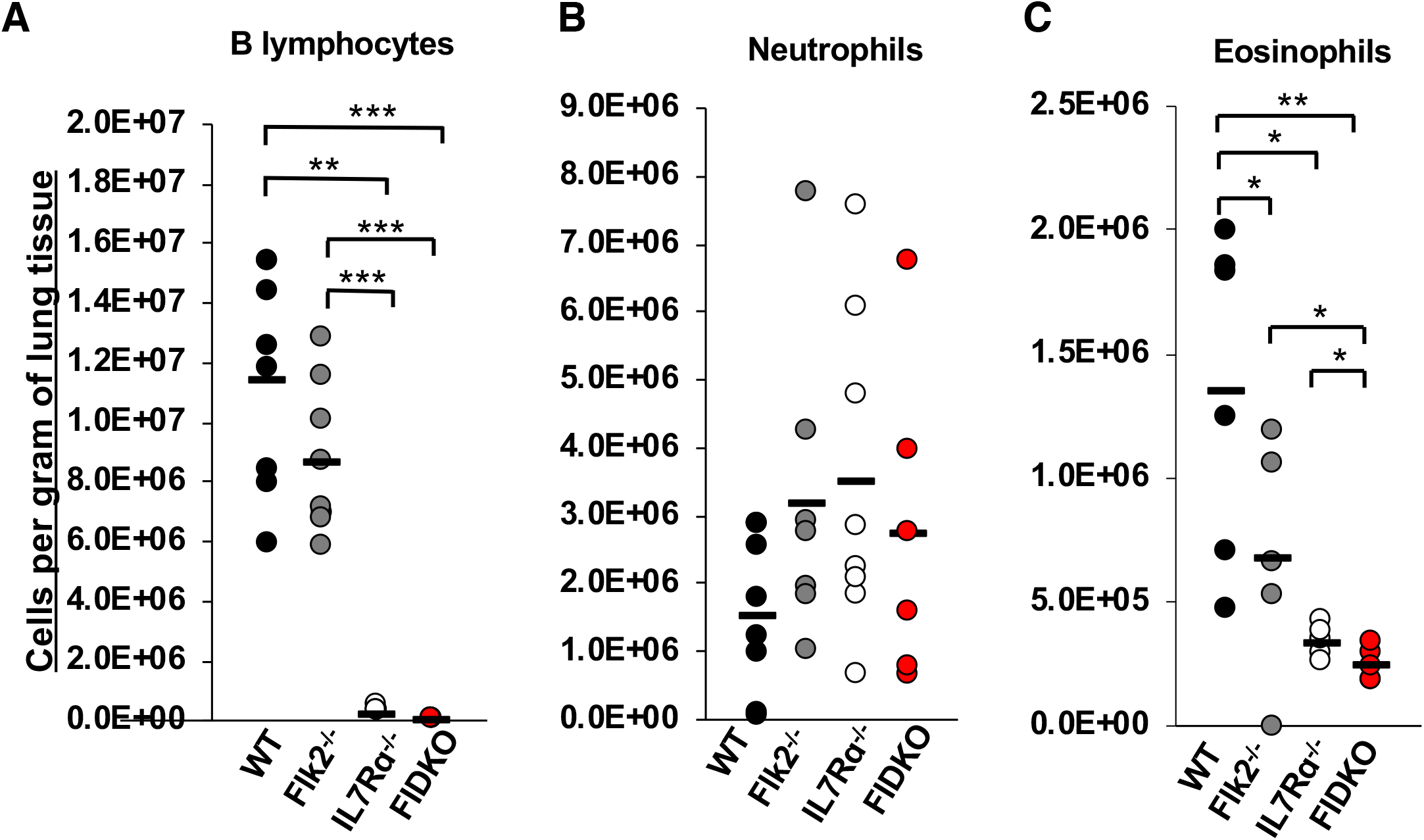
Deletion of Flk2 and/or IL7Rα differentially affects numbers of both lymphoid and myeloid cells in the lungs of adult mice. Cellularity was calculated by the total numbers of cells collected and dividing by the weight of the tissue. Dots represent individual mice. Black bars represent the average. *P<0.05, **P<0.005, ***P<0.001. **A**, B lymphocytes were significantly reduced in Il7Rα^-/-^and FIDKO, but not Flk2^-/-^, lungs. Quantification of lymphocytes (Live, Ter119-Mac1-Gr1-CD19+) per gram of tissue in the lungs of wild type (WT) (black), Flk2^-/-^ (gray), IL7Rα^-/-^(white), or FIDKO (red) adult mouse lungs. WT n=8, Flk2^-/-^ n=8, IL7R^-/-^ n=8, and FIDKO n=5 representing four independent experiments. **B**, Neutrophils were not affected by loss of Flk2 or IL7Rα. Quantification of neutrophils (Live, CD3-CD4-CD5-CD8-B220-Ter119-CD45+Ly6g+CD11b+) per gram of tissue in the lungs of wild type (WT) (black), Flk2^-/-^ (gray), IL7Rα^-/-^(white), or FIDKO (red) adult mouse lungs. WT n=8, Flk2^-/-^ n=7, IL7R^-/-^ n=8, and FIDKO n=6 representing four independent experiments. **C**, Eosinophils were significantly reduced in Flk2^-/-^, IL7Rα^-/-^, and FIDKO mouse lungs. Quantification of eosinophils (Live,CD3-CD4-CD5-CD8-B220-Ter119-CD45+Ly6g-CD11b+SiglecF^mid^CD11c-) per gram of tissue in the lungs of wild type (WT) (black), Flk2^-/-^ (gray), IL7Rα^-/-^ (white), or Flk2^-/-^IL7Rα^-/-^ (FIDKO) (red) adult mouse lungs. WT n=6, Flk2^-/-^ n=5, IL7R^-/-^ n=5, and FIDKO n=5 representing four independent experiments.

Surprisingly, deletion of IL7Rα also led to a significant reduction in numbers of eosinophils in the lungs, with a further significant reduction in the FIDKO mice (Fig. 1C). To determine whether this was a tissue-specific phenotype, we also analyzed the PB and BM of WT and IL7R^-/-^ mice. We observed that eosinophils were similarly affected in these tissues (Fig. S1B and D). Interestingly, there was no significant difference in neutrophils numbers in the BM of these mice, but there were significantly more circulating neutrophils in the IL7Rα^-/-^ mice (Fig. S1A and C). These data indicate that IL7R is differentially required for eosinophil and neutrophil homeostasis across tissues.

### IL7R-Cre does not label adult neutrophils or eosinophils

Recently, we established that IL7R-Cre, but not Flk2-Cre, robustly labels trMacs across several tissues due to transient, developmental expression of IL7Rα in the monocyte-macrophage lineage^6^. To interrogate whether eosinophils have a history of IL7R expression, we crossed mice expressing IL7R-Cre^16^ to mTmG mice containing a dual color fluorescent reporter^17^, thereby creating the “IL7RαSwitch” model analogous to the previously described FlkSwitch mouse (Fig. 2A-2B)^6,13,14,18^. In both models, all cells express Tomato (Tom) unless Cre-mediated recombination cause an irreversible switch to GFP expression by Cre-expressing cells and all of their progeny (Fig. 2A-2B). Labeling of lung B lymphocytes was high (>95%) in both models (Fig. 2C), as has previously been reported for circulatory lymphocytes^6,16^. Similar to myeloid cells in the peripheral blood (PB), labeling of lung neutrophils and eosinophils was high (>95%) in the FlkSwitch mice (Fig. 2D-2E). In contrast, in the IL7RαSwitch mice labeling of lung neutrophils was nominal (<3%) (Fig. 2D), similar to previously reported PB neutrophils^6,16^. Despite the significant reduction in eosinophil numbers in IL7Rα^-/-^ mice (Fig. 1C), labeling of eosinophils was also nominal (<3%) in the IL7RαSwitch mice (Fig. 2E). These data show that most lung eosinophils do not develop through an IL7Rα+ pathway yet depend on IL7R for maintenance of normal cell numbers.

**Figure 2.**
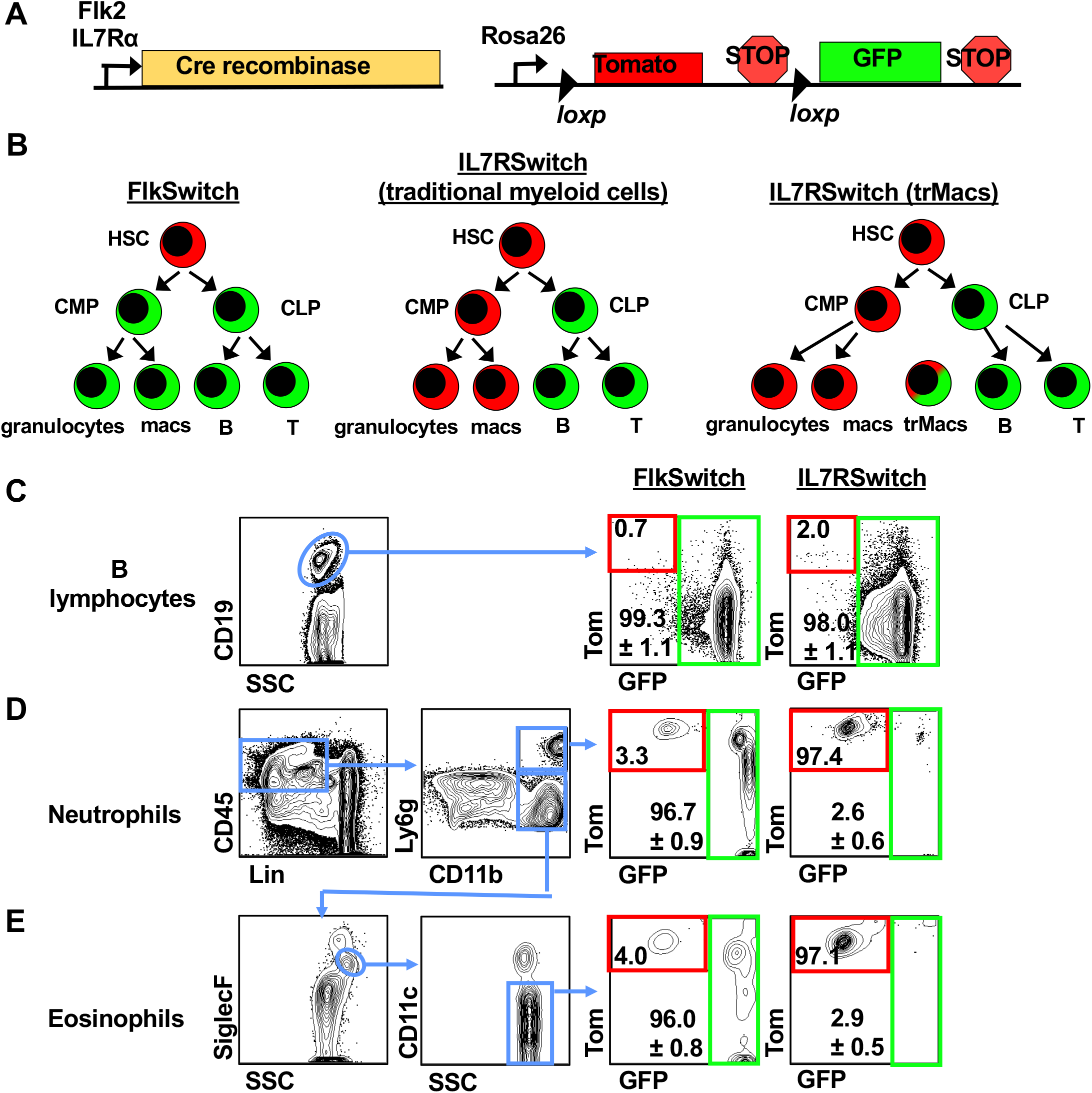
IL7R-Cre efficiently labels lung B cells but does not label adult lung neutrophils or eosinophils. **A**, Schematic of the ‘Switch’ models. Cre recombinase expression was controlled by either Flk2 or IL7Rα regulatory elements. Cre-driver mice were crossed to the mTmG mice expressing a dual-color reporter expressing either Tom or GFP, under control of the Rosa26 locus. Expression of Cre results in an irreversible genetic deletion event that causes a switch in reporter expression from Tom to GFP. **B**, Schematic of Cre-mediated reporter switching in the ‘switch’ models. All cells express Tom unless expression of Cre resulted in an irreversible switch from Tomato to GFP expression. Once a cell expresses GFP, it can only give rise to GFP-expressing progeny. **C**, Lung B cells were highly labeled by both Flk2-Cre and IL7R-Cre lineage tracing. Representative flow cytometric analysis of reporter expression in CD19+ lymphocytes in the lungs of adult FlkSwitch and IL7RSwitch mice. Tomato (Tom) and GFP expression is highlighted by red and green boxes, respectively, in FlkSwitch and IL7Rswitch models. **D**, Lung neutrophils were highly labeled by Flk2-Cre, but not IL7R-Cre, lineage tracing. Representative flow cytometric analysis of reporter expression in CD45+Lin-Ly6g^hi^ neutrophils in the lungs of adult FlkSwitch and IL7Rswitch mice. Tom/GFP gating strategies as in panel C. **E**, Lung eosinophils were highly labeled by Flk2-Cre, but not IL7R-Cre, lineage tracing. Representative flow cytometric analysis of reporter expression in CD45+Lin-Ly6g^lo^SiglecF^mid^CD11c^-^ eosinophils in the lungs of adult FlkSwitch and IL7Rswitch mice. Plots and values are representative of at least three or four mice per cohort from four independent experiments. Values indicate mean frequencies±S.E.M. of gated GFP+ populations.

### IL7Rα^-/-^ HSCs efficiently generate lung eosinophils upon transplantation

To determine whether IL7R-deficient progenitors lack the intrinsic potential to make eosinophils, we transplanted IL7Rα^-/-^ or wild type (WT) HSCs into sublethally irradiated WT recipients and analyzed immune cell reconstitution in the blood and lung >4 months post-transplantation (Fig. 3A). As expected, the IL7Rα^-/-^ HSCs showed diminished ability to produce circulating lymphoid, but not “traditional” myeloid, cells in the blood of the same mice (Fig. S2A,B). In contrast, lung neutrophils and eosinophils were generated with equal efficiency by WT and IL7Rα^-/-^ HSCs (Fig. 3B). The strong reconstitution potential of eosinophils and neutrophils of the lung, comparable to the GM donor chimerism in the periphery, further supports that these cells are “traditional”, adult HSC-derived myeloid cell types and that they do not intrinsically require IL7R for their development.

**Figure 3:**
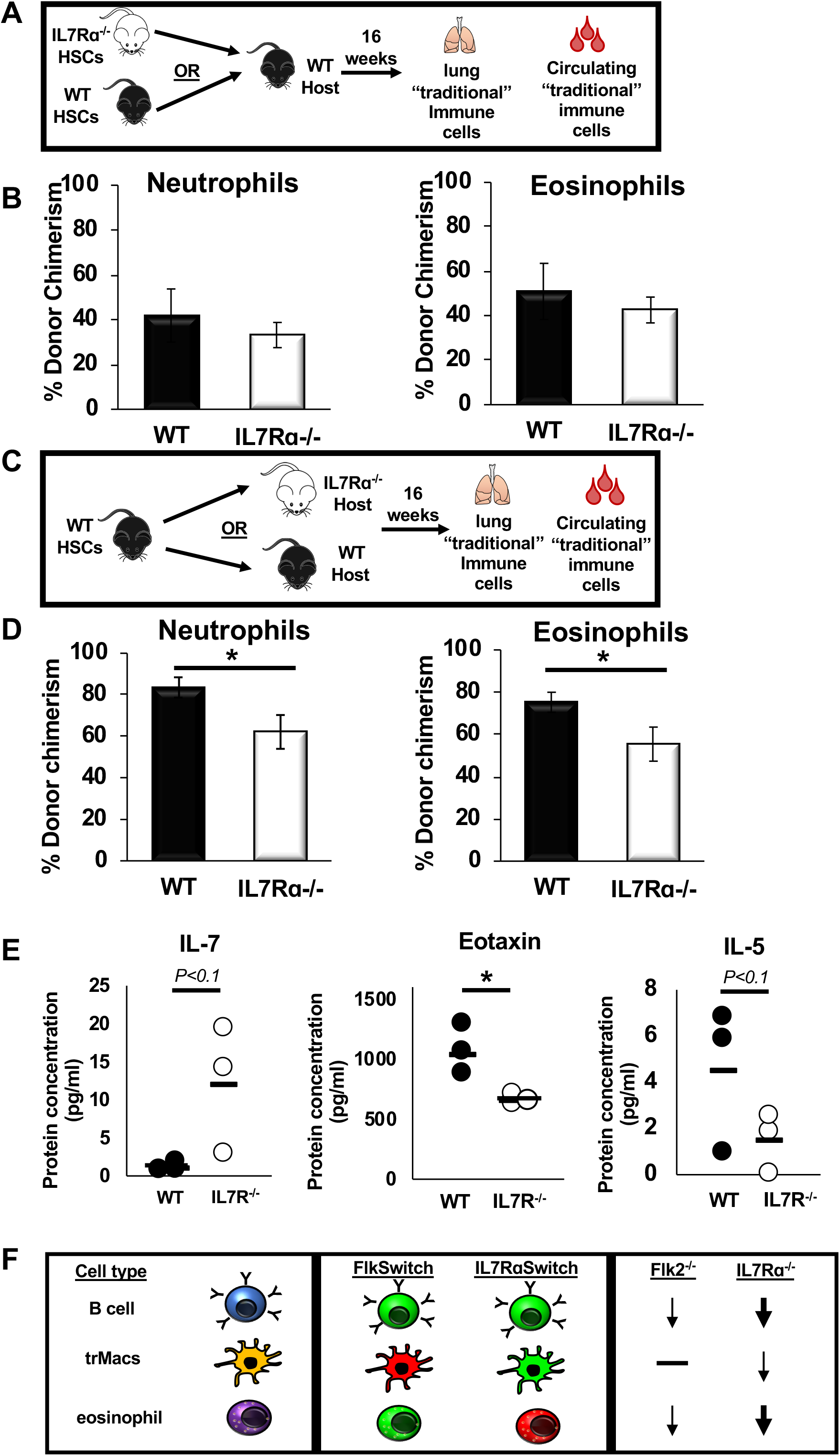
IL7Rα is extrinsically required for eosinophil homeostasis in the lung. **A**, Schematic depicting the transplantation experimental setup used to determine whether IL7R regulates eosinophil development by cell intrinsic mechanisms. 500 WT or IL7Rα^-/-^ HSCs were transplanted into a ¾ sublethally irradiated wild type (WT) GFP recipient. After 16 weeks post transplant, the lungs were harvested and mature immune cells were analyzed via flow cytometry for donor chimerism. **B**, IL7R deletion did not alter the ability of HSCs to reconstitute lung eosinophils. Percent donor chimerism of neutrophils and eosinophils in the lungs of transplanted mice. Error bars are SEM, WT n = 4, IL7Rα^-/-^ n=6 from 2 independent experiments. **C**, Schematic depicting the transplantation experimental setup used to determine whether cell extrinsic IL7R expression is necessary for eosinophil development and homeostasis. **D**, WT HSCs were less efficient at generating “traditional” myeloid cells in an IL7Rα^-/-^ host. Percent donor chimerism of neutrophils and eosinophils in the lungs of transplanted mice. Error bars are SEM, WT n = 5, IL7Rα^-/-^ n=13 from 4 independent experiments. *P<0.05, **P<0.005, ***P<0.001. **E**, Loss of IL7R results in altered cytokine profile in adult mice. Dot plots representing the protein concentration (in pg/ml) of WT (black dots) or IL7Rα^-/-^ (white dots) in the serum of adult mice. Black bars represent the average. WT n=3, IL7Rα^-/-^ n=3 from 3 independent mice; this was sufficient to reach a statistical power of 80% and * P<0.05. **F**, Flk2 and IL7R are differentially involved in the development of multiple hematopoietic cell types. Schematic depicting the differential expression and functional requirement for IL7R and Flk2 in traditional lymphocytes (B cells; top), tissue-resident macrophages (trMacs; middle), and adult lung myeloid cells (eosinophils; bottom). Lymphocytes are highly labeled by both Flk2-Cre (FlkSwitch) and IL7R-Cre (IL7RαSwitch), and functionally dependent on both receptors, although more drastic reductions in cell numbers were observed in IL7Rα^-/-^ (thick arrow) than in Flk2^-/-^mice (thin arrow). In contrast, trMacs are highly labeled by IL7R-Cre, but not Flk2-Cre, and functionally rely on IL7R (thin arrow), but not Flk2 (black line), for efficient development. Here, we report that lung eosinophils are highly labeled by Flk2-Cre, but not IL7R-Cre, yet depend on IL7R (thick arrow), but not Flk2 (black line), for efficient maintenance and reconstitution by HSCs.

### Wild type HSCs fail to fully reconstitute eosinophils in IL7Rα^-/-^ recipients

To determine whether the requirement for IL7R was eosinophil-extrinsic, WT HSCs were transplanted into either WT or IL7Rα^-/-^ recipients (Fig. 3C). WT HSCs efficiently contributed to circulating GMs, Bs, and Ts (Fig. S2C,D), but, intriguingly, displayed significantly impaired reconstitution of neutrophils and eosinophils in the lungs of IL7Rα^-/-^ recipients (Fig. 3D). Taken together, these data suggest that extrinsic IL7R is required for homeostasis of neutrophils and eosinophils in the lung, but not for other “traditional” myeloid cells which circulate in the periphery.

### IL7R deletion caused alterations in cytokines involved in myeloid cell homeostasis

Previous literature has reported several factors implicated in eosinophil development, homing, and survival, including Eotaxin^19,20^, IL-5^21–24^, and others^7,8,25^. To determine which factors may be influencing eosinophil development and survival in the IL7R mutant mice, we collected serum from WT and mutant mice and compared the relative concentration of several cytokines. Because IL-7 has previously been reported to be elevated in IL7Rα^-/-^ mice^7^, we first tested IL-7 levels as our positive control. Consistent with previous data, we observed that IL-7 was upregulated several fold in IL7Rα^-/-^ mice, although borderline statistically significant here (∼10-fold, p<0.1; Fig. 3E). Importantly, we also observed that eosinophil-promoting Eotaxin was significantly downregulated in the IL7Rα^-/-^ mice (Fig. 3E). Additionally, IL-5 was decreased 3-fold in the mutant mice, near statistical significance (p<0.1; Fig. 3E). Taken together, these data indicate that downregulation of Eotaxin and IL-5, and possibly additional factors, may play important roles in eosinophil development and survival in the IL7R mutant mice.

Our data presented here reveal that IL7R is required for specific myeloid cell homeostasis in the lungs, PB, and BM of adult mice. Other groups have reported that human eosinophils have detectable levels of IL7Rα mRNA and surface protein^7,8^, and that IL7R mRNA can be induced in human monocytes with LPS stimulation^9^. Similarly, we previously found that trMacs in the lung and other tissues transiently express IL7Rα during development^6^. However, our IL7R-Cre lineage tracing data of traditional lung myeloid cells reported here show that only a small proportion of eosinophils and neutrophils are labeled at steady state. This strongly argues against a requirement for expression by eosinophils or their precursors. Instead, we favor a model of cell-extrinsic requirement for IL7Rα in eosinophil development. This notion is supported by the reciprocal transplantation assays, where we found that WT and IL7Rα^-/-^ HSCs are equally capable of contributing to eosinophils and neutrophils in WT hosts (Fig. 3B). Conversely, when we transplanted WT HSCs into an IL7R null background, we observed that the donor chimerism of eosinophils and neutrophils in the lungs were significantly impaired (Fig. 3D), but that circulating “traditional” myeloid cells were efficiently generated (Fig. S2B). Interestingly, it has been reported that eosinophil homeostasis relies on lymphocyte- and stromal-secreted survival factors^21,22,25–28^. We found that two known eosinophil regulators, Eotaxin and IL-5, were reduced in the IL7Rα^-/-^ mice. Both of these cytokines are known to be secreted by lymphoid cells to promote eosinophil development, homing, and survival^19,20,22–24,29^. Additionally, as previously reported^7^, we observed that IL-7 was upregulated in the IL7R^-/-^ mice, likely due to unbound excess IL-7 in the absence of lymphoid cells. These data could potentially explain the mechanisms behind the eosinophil decrease in IL7Rα^-/-^ mice (Fig. 1C, S1B,D, 3D), which have drastically reduced numbers of lymphocytes^16^ (Fig. 1A, S2B,D). The lack of lymphoid cells results in less secretion of Eotaxin and IL-5, resulting in poor eosinophil development and survival. Taken together, this suggests an extrinsic requirement for IL7R for eosinophil and neutrophil homeostasis. The data reported here represent a novel role of IL7R in “traditional” myeloid cell homeostasis, in a cell type-specific manner. We interpret our findings to suggest a cell-intrinsic role for IL7R in lymphoid cell development and survival, which then, in turn, support traditional myeloid cells in the lungs of adult mice. The results presented here add to the body of knowledge of how complex and dynamic Flk2 and IL7R expression and function cooperate in the regulation of a variety of immune cells, including traditional myeloid cells in the adult lung (Fig. 3F). These findings are significant for lung health, because understanding hematopoietic homeostasis in the lung provides insight on susceptibility to respiratory disease.

## Methods

### Mice

All animals were housed and bred in the AALAC accredited vivarium at UC Santa Cruz and group housed in ventilated cages on a standard 12:12 light cycle. All procedures were approved by the UCSC Institutional Animal Care and Use (IACUC) committees. IL7Rα-Cre^6,16^ and Flk2-Cre (Benz et al 2008) mice, obtained under fully executed Material Transfer Agreements, were crossed to homozygous Rosa26^mTmG^ females^17^ to generate “switch” lines, all on the C57Bl/6 background. WT C56Bl/6 mice were used for controls and for all expression experiments. Adult male and female mice were used randomly and indiscriminately, with the exception of FlkSwitch mice, in which only males were used because of more uniformly high floxing in male than in female mice.

### Tissue and cell isolation

Mice were sacrificed by CO_2_ inhalation. Adult peripheral blood was collected by femoral artery knick. Lungs were dissected, manually dissociated, and incubated in 1x PBS(+/+) with 2% serum, 1-2mg/ml collagenase IV (Gibco) with 100U/ml Dnase1 for 1-2 hours. Following incubation, all tissues were passed through a 16g needle ∼10X followed by a 19g needle ∼10X to make a single cell suspension, and then filtered through a 70 µM filter to obtain a single cell suspension. Cells were pelleted by centrifugation (1200g/ 4 degrees C /5 minutes). Numbers neutrophils, eosinophils, and B lymphocytes were analyzed and compared from the same tissue preparations from the same mice.

### Flow Cytometry

Cell labeling was performed on ice in 1X PBS with 5 mM EDTA and 2% serum^6^. Analysis was performed on a customized BD FACS Aria II and analyzed using FlowJo. Antibodies used: cd45.2-PB (BioLegend-109820), Ter119-PGP (BioLegend-116202), CD3-PGP (BioLegend-100202), CD4-PGP (BioLegend-100402), CD5-PGP (BioLegend-100602), CD8-PGP (BioLegend-100702), B220-PGP (BioLegend-103202), CD11b-PGP (BioLegend-101202), Gr1-PGP (BioLegend-108402), Ly6g-APC (BiolLegend-127614), CD11b-PeCy7 (BiolLegend-101216), SiglecF-BV786 (BioLegend-740956), CD11c-APC-Cy7 (BioLegend-117323), B220-APC Cy7 (BioLegend-103224), CD45.2-A700 (Biolegend-109822), CD19-BV786 (Biolegend-115543), GAR-PE Cy5 (Life Technologies-A-10691).

### Transplantation Assays

Transplantation assays were performed as previously described^11,18,30–34^. Briefly, sorted cells were isolated from wild type (WT) or IL7Rα knockout mice (IL7Rα^-/-^) donor bone marrow. WT recipient mice aged 8-12 weeks were sublethally irradiated (750 rad, single dose) using a Faxitron CP-160 (Faxitron). Under isofluorane-induced general anesthesia, sorted cells were transplanted retro-orbitally.

### Cytokine Analysis

Serum was harvested from wild type or IL7R^-/-^ mice via cardiac puncture. Several hundred microliters of blood was collected and transferred to a 1.5ml anti-coagulant free tube and allowed to clot at room temp for 30 min. Blood was centrifuged at 3,000 rpm for 10 min at 4C. Serum was carefully transferred to a fresh tube and stored at -80C until shipment to Eve Technologies (Calgary, Canada) for analysis..

### Quantification and Statistical Analysis

Number of experiments, n, and what n represents can be found in the legend for each figure. Statistical significance was determined by two-tailed unpaired student’s T-test. All data are shown as mean ± standard error of the mean (SEM) representing at least two to three independent experiments. Power calculations for cytokine analysis indicated that 3 mice was sufficient to achieve a statistical power of 80% at 0.9-0.95 confidence level (https://epitools.ausvet.com.au/twomeanstwo).

## Acknowledgments

We thank Dr. I. Lemischka for the Flk2^-/-^ mice; Drs. H-R. Rodewald and S.M. Schlenner for the IL7Rα-Cre strain; Dr. T. Boehm for the Flt3^Cre^ strain; Bari Nazario and the UCSC Institute for the Biology of Stem Cells for flow cytometry support.

## Author contributions

T.C., A.W. and E.C.F. conceived of the study, designed the experiments, and co-wrote the paper. T.C., A.W., D.P., and A.H. performed experiments and analyzed data. All authors reviewed the manuscript.

